# Cell type transcriptional identities are maintained in cultured *ex vivo* human brain tissue

**DOI:** 10.1101/2024.12.19.629223

**Authors:** JP McGinnis, Joshua Ortiz-Guzman, Sai Mallannagari, Maria Camila Guevara, Benjamin D. W. Belfort, Suyang Bao, Snigdha Srivastava, Maria Morkas, Emily Ji, Kalman A. Katlowitz, Angela Addison, Evelyne K. Tantry, Melissa M. Blessing, Carrie A. Mohila, Nisha Gadgil, Samuel G. McClugage, David F. Bauer, William E. Whitehead, Guillermo Aldave, Omar Tanweer, Naser Jaleel, Ali Jalali, Akash J. Patel, Sameer A. Sheth, Howard L. Weiner, Shankar Gopinath, Ganesh Rao, Akdes Serin Harmanci, Daniel Curry, Benjamin R. Arenkiel

**Author notes:** These authors contributed equally (JP McGinnis, Joshua Ortiz-Guzman). These authors contributed equally (Sai Mallannagari, Maria Camila Guevara).

## Abstract

It is becoming more broadly accepted that human-based models are needed to better understand the complexities of the human nervous system and its diseases. The recently developed human brain organotypic culture model is one highly promising model that requires the involvement of neurosurgeons and neurosurgical patients. Studies have investigated the electrophysiological properties of neurons in such *ex vivo* human tissues, but the maintenance of other cell types within explanted brain remains largely unknown. Here, using single-nucleus RNA sequencing, we systematically evaluate the transcriptional identities of the various cell types found in six patient samples after fourteen days in culture (83,501 nuclei from day 0 samples and 45,738 nuclei from day 14 samples). We used two pediatric temporal lobectomy samples, an adult frontal cortex sample, two IDH wild-type glioblastoma samples, and one medulloblastoma sample. We found remarkably high correlations of day 14 transcriptional identities to day 0 tissue, especially in tumor cells (r = 0.90 to 0.93), though microglia (r = 0.86), oligodendrocytes (r = 0.80), pericytes (r = 0.77), endothelial cells (r = 0.78), and fibroblasts (r = 0.76) showed strong preservation of their transcriptional profiles as well. Astrocytes and excitatory neurons showed more moderate preservation (r = 0.66 and 0.47, respectively). Because the main difficulty with organotypic brain cultures is the acquisition of human tissue, which is readily available to neurosurgeons, this model is easily accessible to neurosurgeon-scientists and neurosurgeons affiliated with research laboratories. Broad uptake of this more representative model should prompt advances in our understanding of many uniquely human diseases, lead to more reliable clinical trial performance, and ultimately yield better therapies for our patients.

## Introduction

The translational limitations of animal models are widely acknowledged and contribute to the lack of meaningful advances for some of our most challenging neurological diseases^1,2^. The need for human-based models with greater predictive validity for human biology, especially the nervous system, is becoming more widely accepted (see, e.g., the NIH Common Fund’s new Complement-ARIE program). One example of this more human-centric approach is cerebral organoid work. However, current organoid protocols at best mimic human neurons no more mature than second trimester fetal brains^3^. More relevant to neurosurgeons and typical neurosurgical diseases is the recently developed human brain organotypic slice model, which maintains human brain tissue in a living state *ex vivo*^4^. To serve as reliable predictors of human nervous system responses to experimental interventions and therapeutic screens, *ex vivo* brain tissue would ideally maintain the relevant features of the brain’s cell types across a reasonable time in culture.

Accordingly, several groups have studied the ability of organotypic cultures to preserve the electrophysiological profiles of human neurons^5,6,7,8,9,10^. In order to examine the wider array of cells found within the brain, especially in such pathologies as tumors, here we used single-nucleus RNA sequencing to profile the stability of individual cell types over fourteen days in culture. Our goal was to establish a baseline for how well different cell type identities in *ex vivo* brain tissue were preserved across two weeks in culture, using methods accessible and familiar to neurosurgeons and common to molecular biology labs. The relative preservation of these transcriptional profiles can be used to compare different model systems’ fidelity to human brain tissue, and to evaluate different organotypic culture conditions. By preserving *ex vivo* human brain tissue in a transcriptional state as near to its *in vivo* profile as possible, we can increase the predictive validity of our models, see more reliable clinical trial performance, and more quickly generate useful therapies for our patients^1^.

We have therefore performed, for each of six patient samples, single-nucleus RNA sequencing of tissue frozen shortly after resection (‘day 0’) as well as tissue from the same patient cultured for fourteen days (‘day 14’). The samples included three tissues with more normal cell types (two pediatric temporal lobe cortex samples and one adult frontal cortex sample overlying a deeper insular tumor), two IDH wild-type glioblastoma tumor samples, and one SHH-type medulloblastoma sample. For each patient sample, we generated a list of genes that were significantly up- or down-regulated (adjusted p-value < 0.05) for each day 0 cell type. By doing this, we obtained a list of genes and their relative expression levels that uniquely defined each cell type’s identity just after resection. We then generated a second list of up-and down-regulated genes in the same way but substituted, one at a time, a day 14 cell type for the corresponding day 0 cell type, generating the significantly up- and down-regulated genes that defined the day 14 cell types. We then asked how well a given cell type’s day 14 relative expression profile correlated with the corresponding day 0 cell type’s expression profile for that patient’s sample. By comparing cell types that have been maintained for fourteen days in culture to those frozen just after resection, we evaluated how well cell types maintained their transcriptional identities, and by extension, the *in vivo* identities of those cell types in the living human brain.

## Results

Three patient samples were expected to contain relatively normal cell types, though were not strictly normal: a 2-year-old patient’s temporal lobectomy sample (Sturge-Weber syndrome, a neurocutaneous syndrome frequently affected by epilepsy, tissue A), a 6-year-old patient’s temporal lobectomy sample (cortical migrational abnormalities, tissue B), and a 60-year-old patient’s frontal lobe cortex (overlying a deeper insular glioma, tissue C) (Figure 1b).

**Figure 1.**
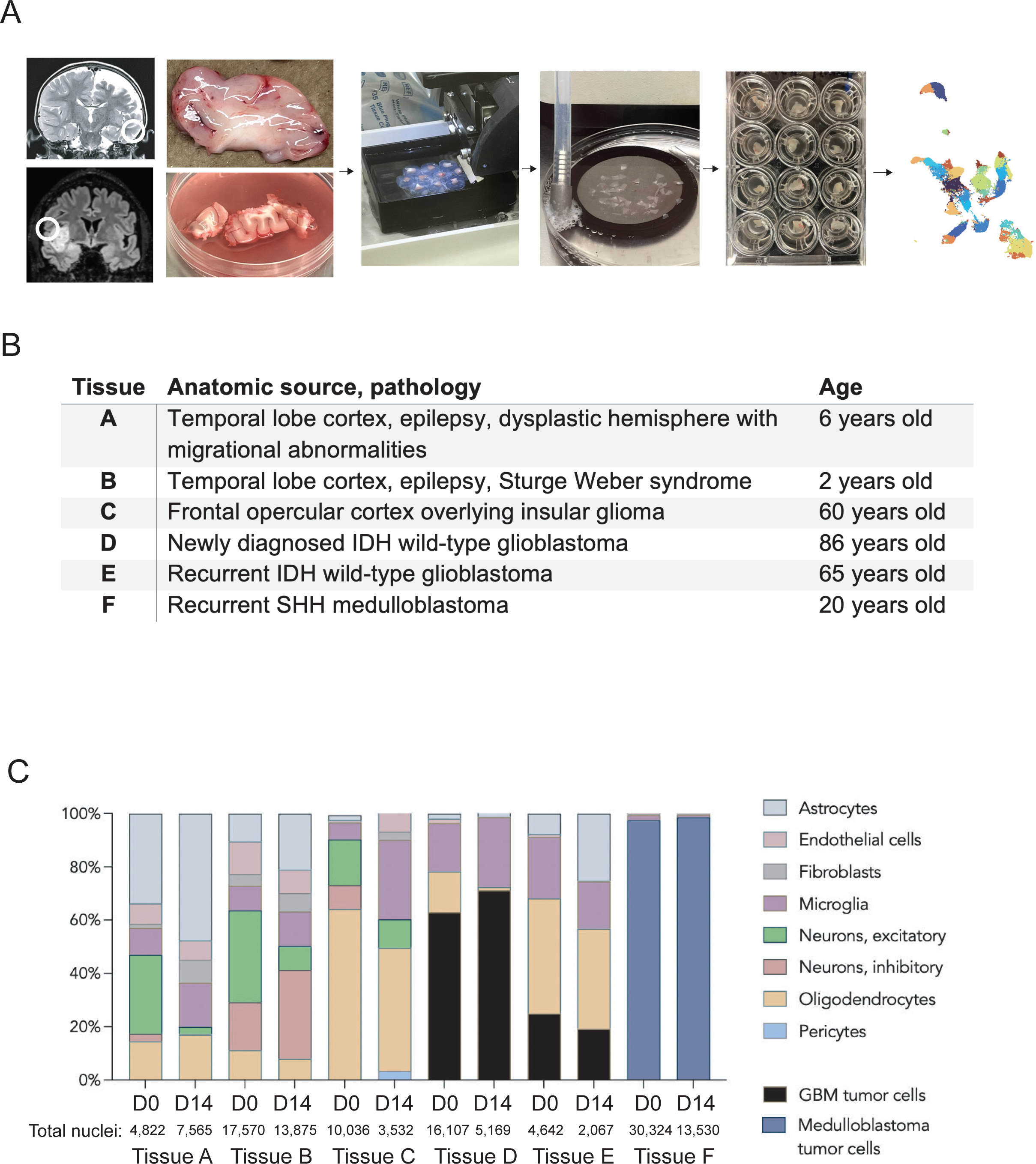
The human brain organotypic slice culture model is feasible and shows conservation of most cell types across time. **A,** Workflow of the human brain organotypic slice culture model, from MRI to tissue sectioning to culture plates to single-nucleus RNA sequencing. **B**, List of tissues used in this study with anatomic locations and pathologies, as well as approximate ages. **C**, Bar graphs showing the relative proportions of identified cell types in day 0 and day 14 samples for each patient tissue. Cell types were identified using canonical marker gene expression from published single-cell and single-nucleus RNA sequencing data. Number of nuclei that passed quality control and were used for analysis listed for each time point.

Two samples were obtained from patients undergoing resection of IDH wild-type glioblastomas—one case newly diagnosed in an 80-year-old (tissue D), the other a 60-year-old patient with recurrent glioblastoma who had undergone a prior surgery and temozolomide/radiation (tissue E). In these cases we used tissue at the enhancing margin or just outside of it, avoiding the necrotic core. The last sample was a recurrent SHH-type medulloblastoma from a 20-year-old patient (tissue F).

### Tissue composition

Once sequenced and aligned to the reference human genome (GRCh 38), each patient sample’s day 0 and day 14 data were integrated and plotted in uniform manifold approximation and projection (UMAP) space. Single-cell RNA sequencing clusters were annotated manually, using combinations of published marker genes (Supplemental Figures 1-6)^11,12^. Once cluster identity was assigned, we assessed the proportions of cell types present in each patient sample at day 0 and day 14 to determine changes in cell type composition over two weeks in culture (Figure 1c). In most tissues, comparing the day 14 back to the day 0 samples, the astrocyte and fibroblast populations increased in relative proportion, while microglia and oligodendrocytes remained generally stable between samples. The share of neurons notably fell over time, and in the adult cortical sample, inhibitory neurons and astrocytes were no longer seen. In tumor samples, the share of malignant cells remained largely stable across fourteen days in culture (Figure 1c).

### Measures of gene expression in normal cell types

We integrated the day 0 and day 14 samples for each patient, and using UMAP plots generated for tissues A, B, and C, we asked whether cell types from day 14 would be similar enough to day 0 that common cell types would cluster together. We observed that cell types did cluster together almost universally across samples (Figure 2a). We then split the day 0 and day 14 cell to more closely examine each individual cell types within each UMAP (Figure 2b). The one case where clustering was notably different was seen in the 60-year-old frontal lobe cortex (2b, right panel), where day 14 excitatory neurons (right side) clustered separately from the day 0 excitatory neurons (middle). Further, while a small number of astrocytes were present at day 0 (left), none were apparent at day 14.

**Figure 2.**
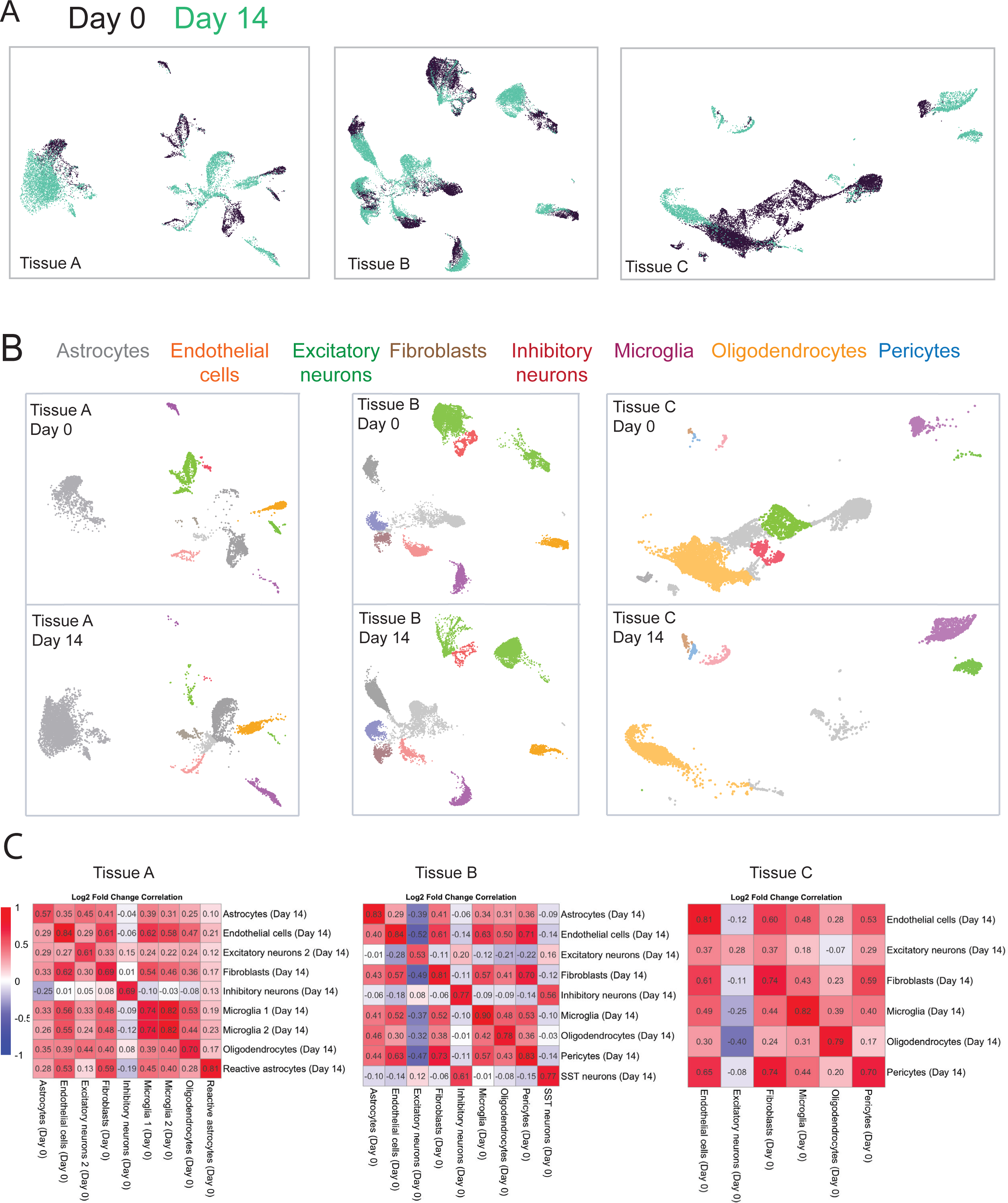
Single-nucleus RNA sequencing shows retained cell type transcriptional identities in tissues with more normal cell types. **A**, Integrated UMAP projections of Day 0 and Day 14 cells for tissues A, B, and C. Each point represents a single cell. **B**, Split UMAPs color-coded by cell type for day 0 and day 14 cells for tissues A, B, and C. Day 0 and day 14 cell types generally cluster together though there are some differences, e.g. tissue C’s neurons in day 0 and day 14. Cell types assigned using published marker genes (see methods). **C**, Correlation matrices comparing the log2 fold change of a given cell type at day 14 to cell types at day 0. Log2 fold changes generated by comparing a given cell type at day 0 to the rest of the day 0 cells, and given cell types at day 14 to the rest of the day 14 cells. Higher correlations at intersections of cell types indicate that the unique transcriptional profile differentiating that cell type from other cell types at day 0 is better preserved at day 14.

We next evaluated whether the distinctive gene expression profile of each cell type was maintained over fourteen days in culture. This was done by first generating a list of all significantly up- or down-regulated genes between cell types in the day 0 sample (adjusted p-value < 0.05). This established the gene expression signatures for each day 0 cell type. We then repeated that analysis but swapped in a given day 14 cell type in place of its day 0 counterpart, generating its signature compared to that patient’s day 0 cell types (and repeated this for each day 14 cell type). In this way we had, for each day 0 and day 14 cell type, a shared list of genes and to what extent a cell type up- or down-regulated the genes on that list. Using this list of genes, we then calculated the correlation coefficients between day 0 and day 14 cell types, in essence asking whether genes that tended to be up- or down-regulated at day 0 also tended to be up- or down-regulated in that same cell type after two weeks in culture. A higher correlation indicates that the unique gene expression profile for a given cell type, the profile that differentiated it from other cell types, was better maintained over two weeks in culture.

Astrocytes showed markedly variable correlations, r = 0.57 and 0.82 in the two samples containing astrocytes at both day 0 and day 14. Endothelial cells showed a consistently higher correlation, r = 0.83, 0.84, and 0.81. Excitatory neurons showed lower and more variable correlations, r = 0.60, 0.52, and 0.28. Fibroblasts (0.69, 0.80, 0.73) and inhibitory neurons when present (0.69, 0.77) showed relatively higher correlations, though not as high as microglia (0.77, 0.89, 0.82). Oligodendrocytes showed moderately high correlation between day 0 and day 14 (0.69, 0.78, and 0.79). Pericytes also showed variability between the two samples where they could be clearly identified (0.83, .70). Overall, all day 14 cell types showed reasonable correlations to their day 0 counterparts, with microglia having the best-preserved gene expression profiles.

### Measures of gene expression in tumors

Because tumor cells may respond differently to *ex vivo* culture conditions, we used two IDH wild-type glioblastoma samples and one medulloblastoma sample to determine whether tumor cell and tumor-associated cell identities would be maintained in culture. We assigned individual cell types to clusters based on published marker genes for the respective tumor types^13,14,15^. Similar to our more normal tissue, day 0 and day 14 cell types clustered together both for glioblastoma and medulloblastoma samples, including the tumor cells themselves (Figure 3a and 3b).

**Figure 3.**
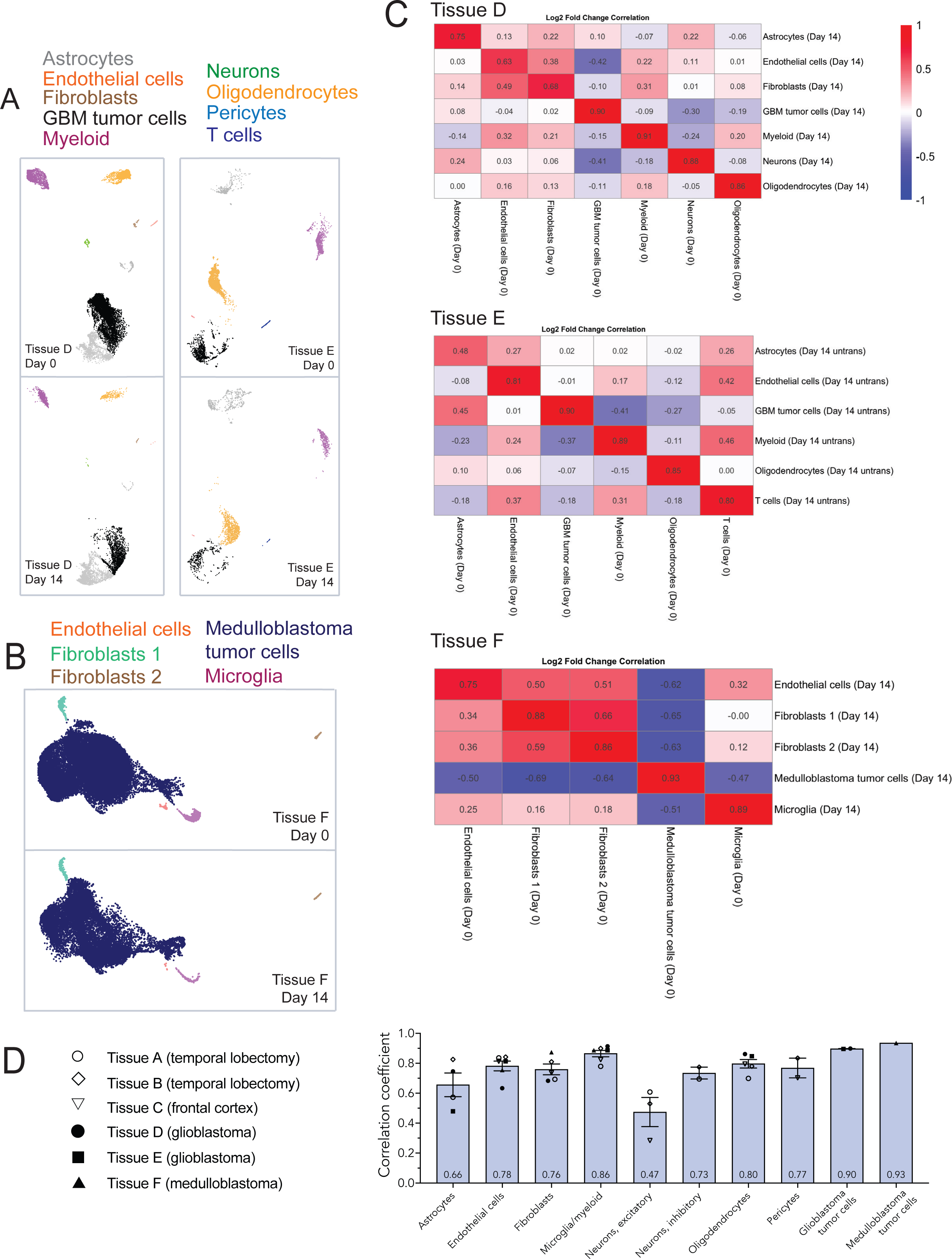
Cells within tumor tissues retain their unique transcriptional profile after two weeks in culture. **A**, Split UMAPs for glioblastoma tissues D and E color-coded by cell types. Some cells were not able to be assigned clearly to a cell type and so left unlabeled (Tissue D, bottom left cluster in lighter grey). **B**, Split UMAP for medulloblastoma tissue (tissue F) color-coded by cell types. **C**, Correlation matrices comparing day 0 to day 14 cell types for tissues D (IDH wild-type glioblastoma), E (IDH wild-type glioblastoma), and F (SHH-type medulloblastoma). **D**, Graph showing the individual values for each tissue sample and the overall average correlation coefficient between day 0 and day 14 cell types. Error bars are standard error of the mean (*n* = 1-6).

Examining the correlation of day 14 cell types to their day 0 counterparts, we found surprisingly high concordance (Figure 3c), especially between tumor cells (tissue D’s GBM tumor cells r = 0.89, tissue E’s GBM tumor cells r = 0.89; medulloblastoma tumor cells r = 0.93). These data indicate that the tumor cells’ gene expression signature is strikingly well-preserved over fourteen days in *ex vivo* culture.

Aggregating common cell types across the more normal and tumor tissues, these data show that the greatest concordance between day 0 and day 14 tissue is seen with tumor cells (medulloblastoma r = 0.93, GBM r = 0.89), followed by microglia (0.86), oligodendrocytes (0.80), endothelial cells (0.78), pericytes (0.77), fibroblasts (0.76), and inhibitory neurons (0.73) (Figure 3d). Astrocytes (0.66) and excitatory neurons (0.47) showed more moderate correlations in transcriptional profiles.

### Differential gene expression between day 0 and day 14 tissue

To complement the assessment of similarity between each cell type at day 0 and day 14, we next calculated the differential expression between each cell type pair for each patient’s day 0 and day 14 samples. We focused on genes differentially expressed in the same direction across at least two patient samples (Supplemental Table 1 for the list of up- and down-regulated genes and in which samples they were identified). We used the established Metascape program to determine whether we could identify any consistent up- or down-regulated gene expression programs across cell types or samples^16^. However, we could not identify any coherent programs that were up- or down-regulated for either non-neoplastic (Supplemental Figure 7) or neoplastic cell types (Supplemental Figure 8), suggesting no clear systemic alteration of these cells’ transcriptional programs while in culture.

## Discussion

We provide here the first set of data using single-nucleus RNA sequencing to systematically compare the fidelity of gene expression profiles in *ex vivo* cultured human brain tissue. Across these six samples, comprising 83,501 nuclei from day 0 samples and 45,738 nuclei from day 14 samples, tumor cells (both glioblastoma and medulloblastoma tumor cells), microglia, oligodendrocytes, endothelial cells, pericytes, fibroblasts, and inhibitory neurons strongly maintained their transcriptional profiles over two weeks in culture (Figure 3d), while astrocytes and excitatory neurons showed moderate preservation.

Our goal was to determine whether gene expression profiles in *ex vivo* human brain tissue are preserved over two weeks in culture, and also establish a baseline so that future improvements in culture conditions can be evaluated by how well they maintain this transcriptional fidelity (in the form of higher correlation coefficients). Second, and more importantly, we think this work demonstrates the feasibility and already high validity of this model. We hope this prompts wider adoption of the model that we think is far more representative of true *in vivo* human biology and pathophysiology than animal models. We anticipate further substantial improvements in culture methods from work being done across several institutions; the decline in neuron number we see here may be mitigated with the use of human CSF as culture media, which is a focus of ongoing work^5^.

The first three samples (tissues A, B, and C) contain more normal cell types, though due to their involvement with epilepsy and an underlying tumor cannot strictly be considered normal. However, future therapeutics that can be tested with this model will not target strictly normal cell types, and so we do not see this as a substantial limitation. It is notable that tumor cells are among the cell types with the greatest transcriptional fidelity to their day 0 counterparts. This conservation is promising for mechanistic studies or tests of interventions that require a certain duration in culture. If human specimens can be kept alive *ex vivo* in a state that predicts a tumor’s response, we should see more reliable clinical trial performance than therapies tested, for example, in xenograft mouse models (or models even less representative). Whether the high correlation seen here holds for other pathologic tissues (such as low-grade gliomas, IDH-mutant high-grade astrocytomas, meningiomas, ependymomas, AT/RTs, DIPG) and rarer cell types is the subject of ongoing study.

That human tissue can maintain its gene expression profile *ex vivo* creates possibilities across a range of fields, from tumors to vascular malformations to epilepsy. For example, manipulating immune cells within human brain tumor tissue, rather than in immunocompromised mice given artificial tumors, could yield important insights that inform future immunotherapies. Manipulating AVM endothelial cell pathways may help guide therapeutic strategies. And finally, using human brain tissue, with its uniquely human cell surface receptors, we can develop targeting modalities capable of cell-type specific transduction, creating therapeutic possibilities for dozens of diseases. By improving the fidelity of our models to *in vivo* human brain, we can drive progress across many areas of neuroscience and neurological diseases.

While this *ex vivo* human brain model is anticipated to have very high predictive validity, there are several limitations, though none insurmountable. First, it relies on a sufficient amount of surgical tissue resected (which depends on the assays required), and sample availability can be unpredictable. Second, it is limited to the types of diseases that are surgically resected or biopsied in sufficient number. One possible avenue for nonsurgical diseases comes from the intriguing finding that early post-mortem tissue can still show synaptic plasticity for several days in culture^17^. Third, the levels of mRNAs are not always the main determinant of protein levels or of a cell’s function, typically the main features of interest^18^. Until the field of single-cell proteomics matures, however, we think single-nucleus (or single-cell) RNA sequencing remains the most useful technique to evaluate simultaneous cell type fidelity across tens of thousands of cells over time. Functional characterization of constituent cell types, beyond electrophysiology, is also needed. Finally, human patient samples should be expected to produce more variability than inbred mouse strains or cell culture lines. This is frequently voiced as a concern, but we and others feel it is a strength of the model, given that clinical trials need to surmount intrinsic human variability^8^.

Because of the vast differences between the human brain and animal models at all levels of analysis, the use of human-based models is vital but currently under-utilized. Not all labs will have ready access to *ex vivo* human brain tissue; those who do have the obligation to collaborate widely to address the most pressing questions that these models can answer. Neurosurgeons have both unique access and the unique clinical perspective that facilitates human brain organotypic studies. Given the relatively low threshold for successfully culturing human brain tissue, and the potential benefits of models with greater predictive validity, we should more broadly adopt and advocate for this model across a wide range of problems and diseases.

## Methods

### Human brain tissue acquisition

Patients planned to undergo resective neurosurgical procedures at three major academic hospitals were approached and consented pre-operatively by a member of the study team (Baylor College of Medicine, IRB H-51865) for any specimens not needed for pathologic diagnosis, with the explicit assurance that participating (or not) would change nothing about their surgery or care. In the case of pediatric patients or adults unable to consent, parents or medical decision-makers were consented. Tissue acquisition was coordinated with the neurosurgery teams at the three hospitals in the Texas Medical Center.

### Human brain tissue culture

Using an IRB-approved protocol, we routinely approach patients scheduled to undergo various neurosurgical resections and obtained consent for any surgical specimens not needed for clinical diagnosis. Specimens were collected from pathology following tissue sectioning and gross examination by board-certified neuropathologists, or directly from the operating room, in sterile plastic specimen containers, usually within minutes and no longer than 45 minutes after resection, into which we poured ice-cold, pre-carbogenated (>20 min bubbling time, 95%/5% O2/CO2) NMDG-aCSF, exactly following the protocol from Park et al. (Park, Nature Protocols 2022). Tissue was rapidly transported to the lab in NMDG-aCSF on ice, where it was manually sectioned inside a biosafety cabinet hood with scalpels into pieces ∼1cm x 1cm. Tissue pieces were placed into a retractable tube and molten ultra-low melting agarose (Sigma A2576) was poured over top. Using a cold clamp the agarose was rapidly solidified, at which point the agarose cylinder was removed and superglued onto a tape-covered vibratome mount. Tissues were oriented so that the sectioning would be done perpendicular to the cortical surface, parallel to neuronal projections. Tissues were sectioned to 300um on a Leica VT1200 vibratome (housed within a biosafety cabinet) in cold aCSF that was continuously bubbled with carbogen. We then moved the slices to room temperature aCSF that was continuously bubbled with carbogen for 15-30 minutes, after which the slices were plated individually onto membranes inserts (CellTreat #230617) overlying 600 µL of organotypic slice culture media (OSCM) whose formulation followed exactly a published OSCM protocol^8^, except that heparin was not added. Any excess media or aCSF was aspirated off the membrane surface, and the plates were kept in a humidified tissue culture incubator at 37 °C, 95% humidity, and 5% CO2. The tissue underwent daily media changes (400 µL removed and added to the surrounding well, avoiding any contact with the tissue slice). At time of collection, slices were gently transferred by forceps into 2 mL eppendorf tubes (2 tissue slices per tube) and flash frozen in liquid nitrogen. Tissues were stored in −80 °C freezers until nuclei isolation.

### Single-nucleus isolation

On the day of processing, 2 to 8 tubes (4 to 16 slices) of frozen tissue were dissociated using GentleMACS nuclei isolation protocol (nuclei isolation buffer [Miltenyi Biotec, cat# 130-128-024], Protector RNAse Inhibitor [Millipore Sigma, cat# 3335402001], GentleMACS C tubes [Miltenyi Biotec, cat# 130-093-237], GentleMACS Octo Dissociator [Miltenyi Biotec, cat# 130-096-427], MACS SmartStrainers 70um [Miltenyi Biotec, cat# 130-098-462], MACS SmartStrainers 30um [Miltenyi Biotec, cat# 130-098-458]). In brief, samples were placed in 2 mL of Miltenyi Nuclei Isolation Buffer and Protector RNAse Inhibitor in GentleMACS C tubes. Samples then underwent the preprogrammed “nuclei isolation” program on a GentleMACS Octo Dissociator. Immediately after dissociation, samples were strained through a 70um MACS SmartStrainer and collected in a 15 ml tube, centrifuged at 300xg for 5 minutes at 4°C. The supernatant was extracted and discarded, and the resulting pellet resuspended in 1mL of ice-cold PBS. Resuspended samples were then run through a 30um MACS SmartStrainer, centrifuged, and resuspended in ∼250-500 µL. Nuclei were then FACS sorted (using DAPI) on a Sony MA900 in the Baylor Cytometry and Cell Sorting Core.

### Single-nucleus RNA library preparation

The single-cell gene expression libraries were prepared by the Baylor Single Cell Core according to the Chromium Single Cell Gene Expression 3’v3.1 instruction (PN-1000121, PN-1000120, PN-1000213, 10x Genomics). Briefly, single cells, reverse transcription (RT) reagents, Gel Beads containing barcoded oligonucleotides, and oil were loaded on a Chromium controller (10x Genomics) to generate single-cell GEMS (Gel Beads-In-Emulsions) where full-length cDNA was synthesized and barcoded for each molecule of mRNA (UMI) and each single cell (cell barcode). Subsequently, the GEMS were broken and cDNA from each single cell was pooled. Following cleanup using Dynabeads MyOne Silane Beads, cDNA is amplified by PCR. The amplified product was fragmented to optimal size before end-repair, A-tailing, and adaptor ligation. The final library was generated by amplification.

### Sequencing

Barcoded and single-nucleus cDNA libraries were submitted for commercial Illumina NovaSeq 2×150 paired-end sequencing (Azenta/Genewiz), targeting ∼1 billion reads per sample, which in the past had given us roughly ∼50,000 reads per cell.

### Cell type annotation

Fastq files were uploaded to the 10X Cloud Analysis web browser and underwent quality control, normalization, and alignment using default parameters. Human genome GRCh38 was used to align the fastq files. Each time point’s data was aligned and normalized separately, and then analyses were integrated using the 10X Cloud server for batch effect correction. Cloupe files were then downloaded from 10X, and clusters were visualized using the 10X Loupe browser. Feature lists were created using published marker genes (Allen Institute Brain Atlas Transcriptomic Explorer; Winkler et al., 2022; Abdelfattah et al., Nature Communications, 2022, and Wang, Acta Neuropathologica Communications, 2023; Karin Ocasio, Nature Communications, 2019) that provided clearer differentiation between cell types (see Supplemental Figures 1-6). UMAP plots were downloaded as SVG files from the 10X Loupe browser.

### Log2 fold change calculation

Using the 10X Loupe browser, we first selected only the day 0 cell types and generated differential expression gene lists for each cell type compared to all other day 0 cells (exact negative binomial test [Yu, Huber, & Vitek, 2013]). We repeated this process for the day 14 cell types, comparing each day 14 annotated cluster with other day 14 clusters for that specific patient sample. We removed genes with minimal expression (average occurrence < 1 count across the dataset), and downloaded CSV files of all significantly variable genes (Benjamini-Hochberg correction, adjusted p-value < 0.05). Custom R code was written to split the CSVs into their individual cell types that contained the gene names, average expression level, log2 fold change, and p-value. The correlation between the log2 fold changes for each cell type were calculated using custom R code. It is important to note that the list of genes contained any gene that was differentially expressed in at least one cell type—the genes across which the log2 fold change was calculated were all significantly variable genes across the patient sample, not just the significantly variable genes for a given cell type. Custom R code used the cell type CSVs to generate the correlation matrices using ggplot. (All code used is immediately available at https://github.com/jpmcginnis/HumanAAVProject and is listed with title “Organotypic paper – “)

Graphs were created in Graphpad Prism 10.

### Differential expression analysis

We used Metascape’s online interface to enter, for each cell type, lists of genes from Table 1 that were upregulated for that cell type, and then again for the list of genes that were downregulated. We only considered genes that were significantly up- or down-regulated in at least two patient samples when available.

### Statistical analysis and rigor

Data are presented as mean ± SEM unless otherwise indicated. For single-nucleus RNA sequencing analysis, differential expression analysis was performed using the 10X Loupe browser algorithm (exact negative binomial test [Yu, Huber, & Vitek, 2013]) with Benjamini-Hochberg correction for multiple comparisons. To be considered significantly variable or differentially expressed, log2 fold changes needed to be less than the adjusted p-value of 0.05.

The six samples used here are the entirety of our samples for which we have day 0 and unperturbed day 14 snRNAseq data, since moving to the air-liquid interface model. We have not excluded any samples.

### Data availability

All code is currently available at https://github.com/jpmcginnis/HumanAAVProject, (named as ‘Organotypic paper -’) and Fastq or Seurat objects are immediately available on request and are pending upload on external-facing servers at Baylor College of Medicine.

## Author contributions

Project design: JPM, JOG, DC, BA

IRB approval: JPM, DC

Human brain tissue consent and acquisition: JPM, JOG, SM, MCG, BB, SS, MM, EJ, AA, ET, MB, CM, NG, SM, DB, WW, GA, OT, NJ, AJ, AP, SS, HW, SG, GR, DC

Tissue processing: JPM, SM, MCG, JOG, EJ, AA, ET Single-nucleus library prep: JPM, MCG, SM, SB

Single-nucleus RNA sequencing analysis: JPM, SM, MCG, SB, ASH

Funding: BA, DC, JOG

Initial manuscript draft: JPM

Manuscript editing and review: all

## Acknowledgements

We thank the hundreds of patients who have generously participated in this project, as well as Hilary, Gemma, Megan, Jennifer, Loretta, Glenn, Omar, Waseem, Johannah, Juan, Melanie, Benssy, and Neal for assistance with tissue acquisition; Mira for assistance with library prep, the Arenkiel lab, cytometry core staff, and the unnamed operating room, pathology, and research staff who have contributed so much time and effort to this work.

## Suppe

**Supplemental Figure 1.**
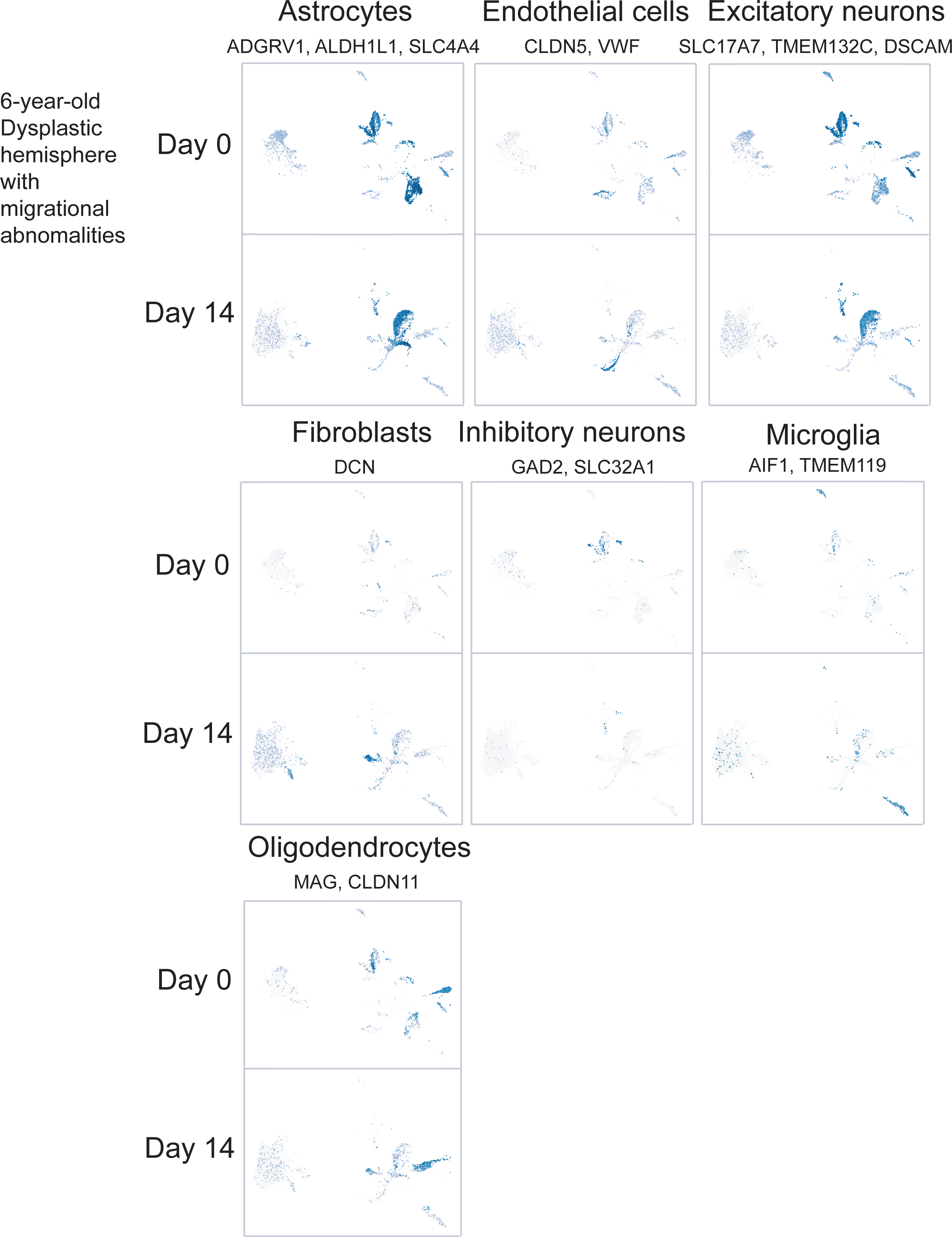
Marker genes used to assign cell types for tissue A. In cases where multiple marker genes were used, feature sum was used across all the genes listed.

**Supplemental Figure 2.**
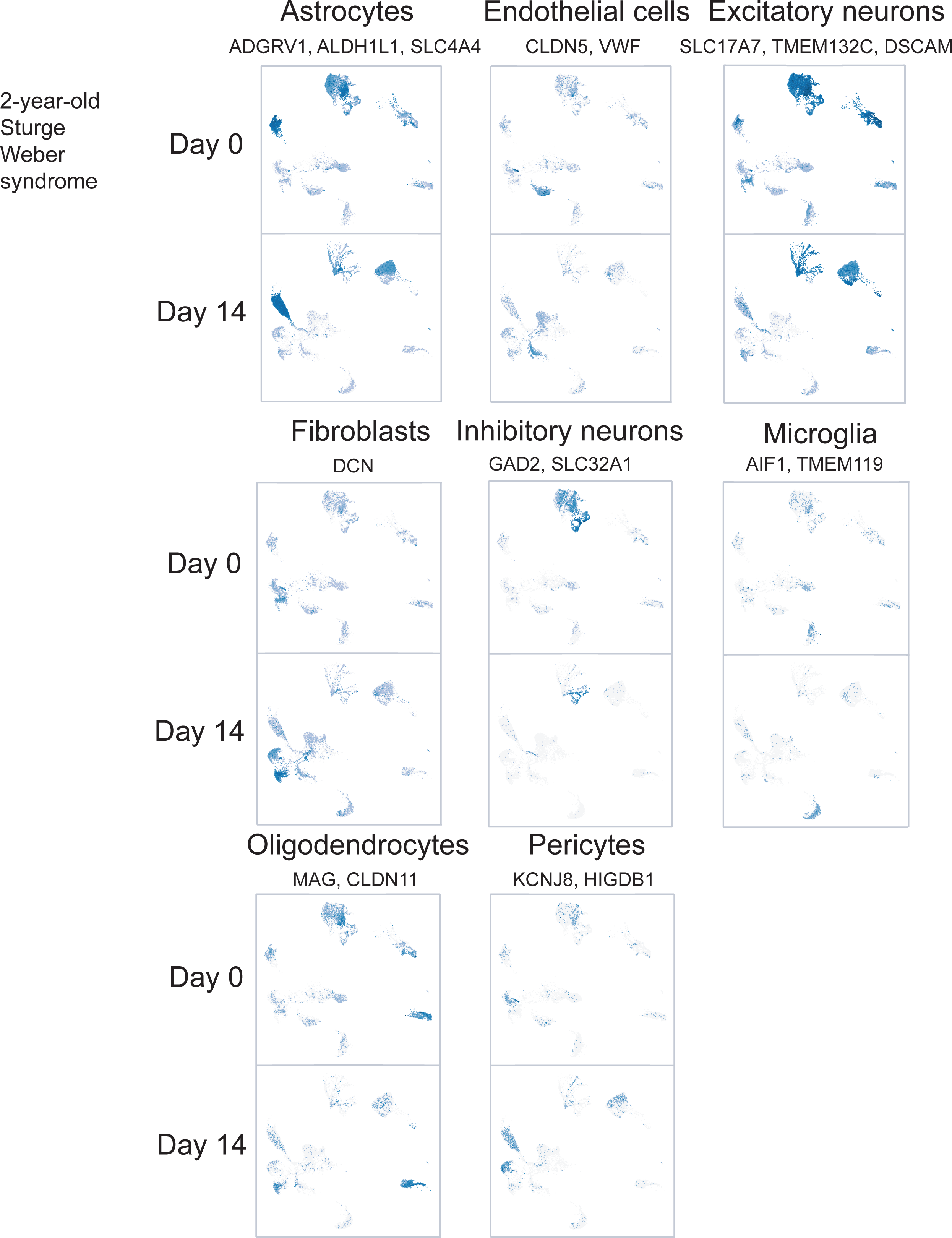
Marker genes used to assign cell types for tissue B. In cases where multiple marker genes were used, feature sum was used across all the genes listed.

**Supplemental Figure 3.**
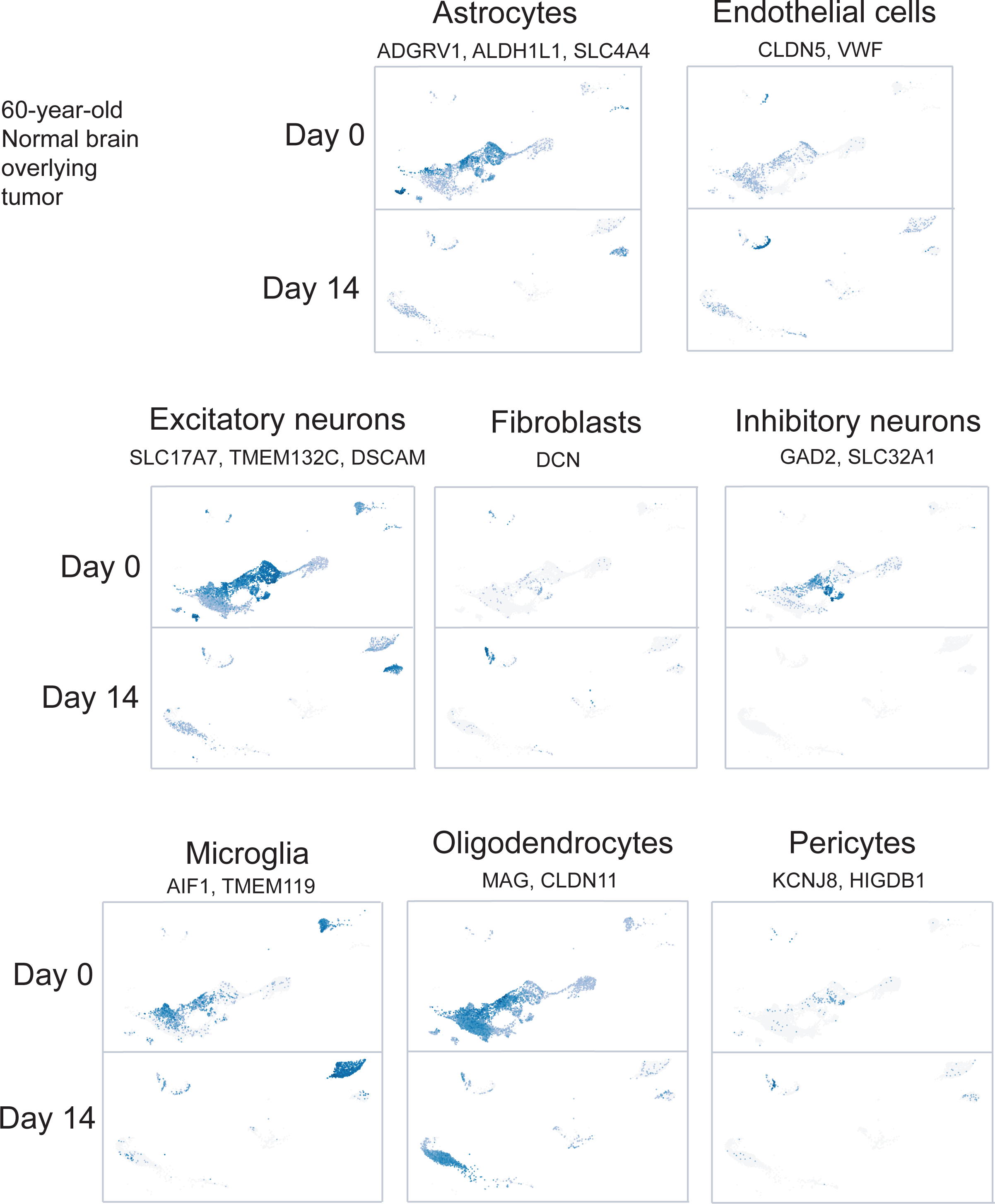
Marker genes used to assign cell types for tissue C. In cases where multiple marker genes were used, feature sum was used across all the genes listed.

**Supplemental Figure 4.**
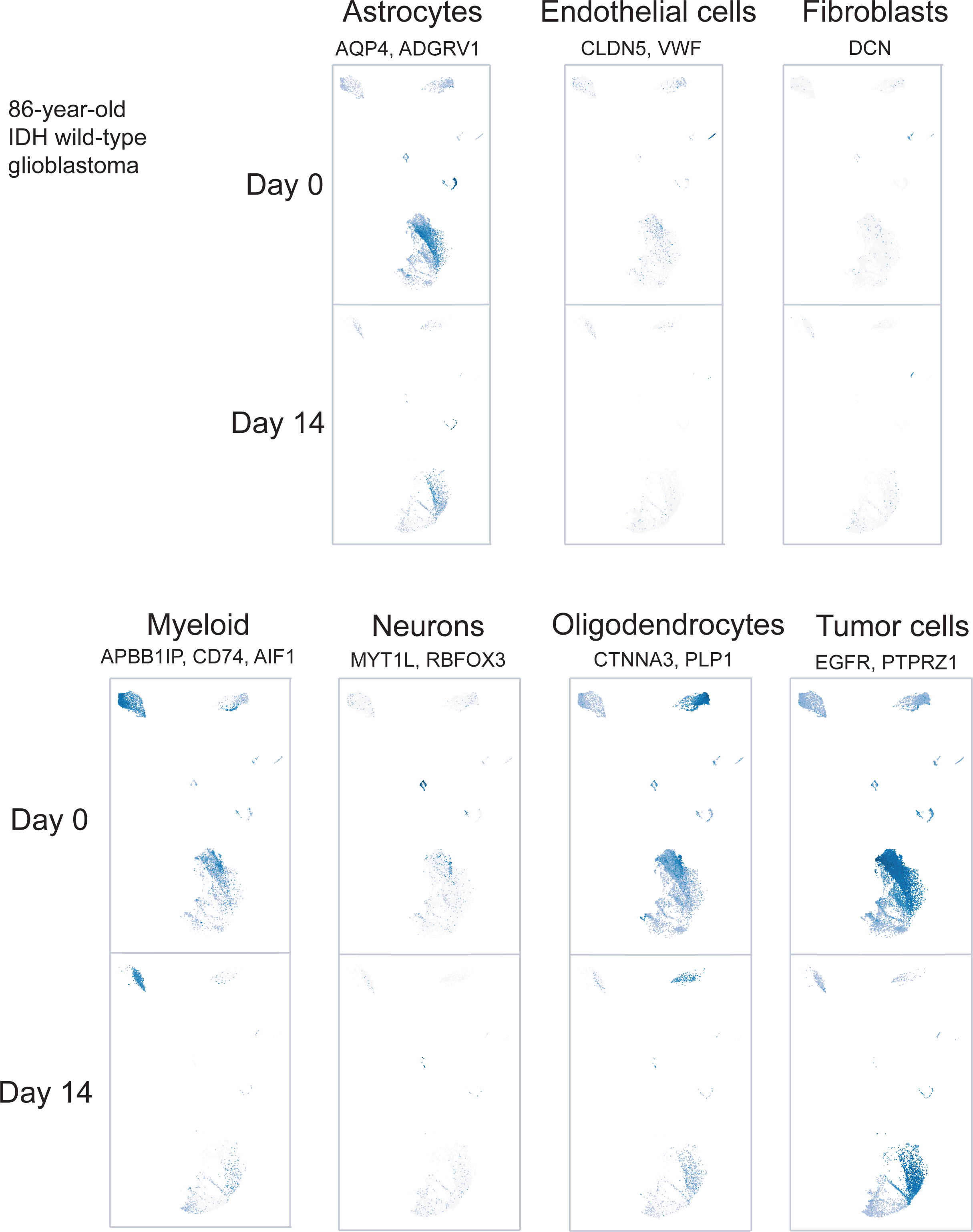
Marker genes used to assign cell types for tissue D. In cases where multiple marker genes were used, feature sum was used across all the genes listed.

**Supplemental Figure 5.**
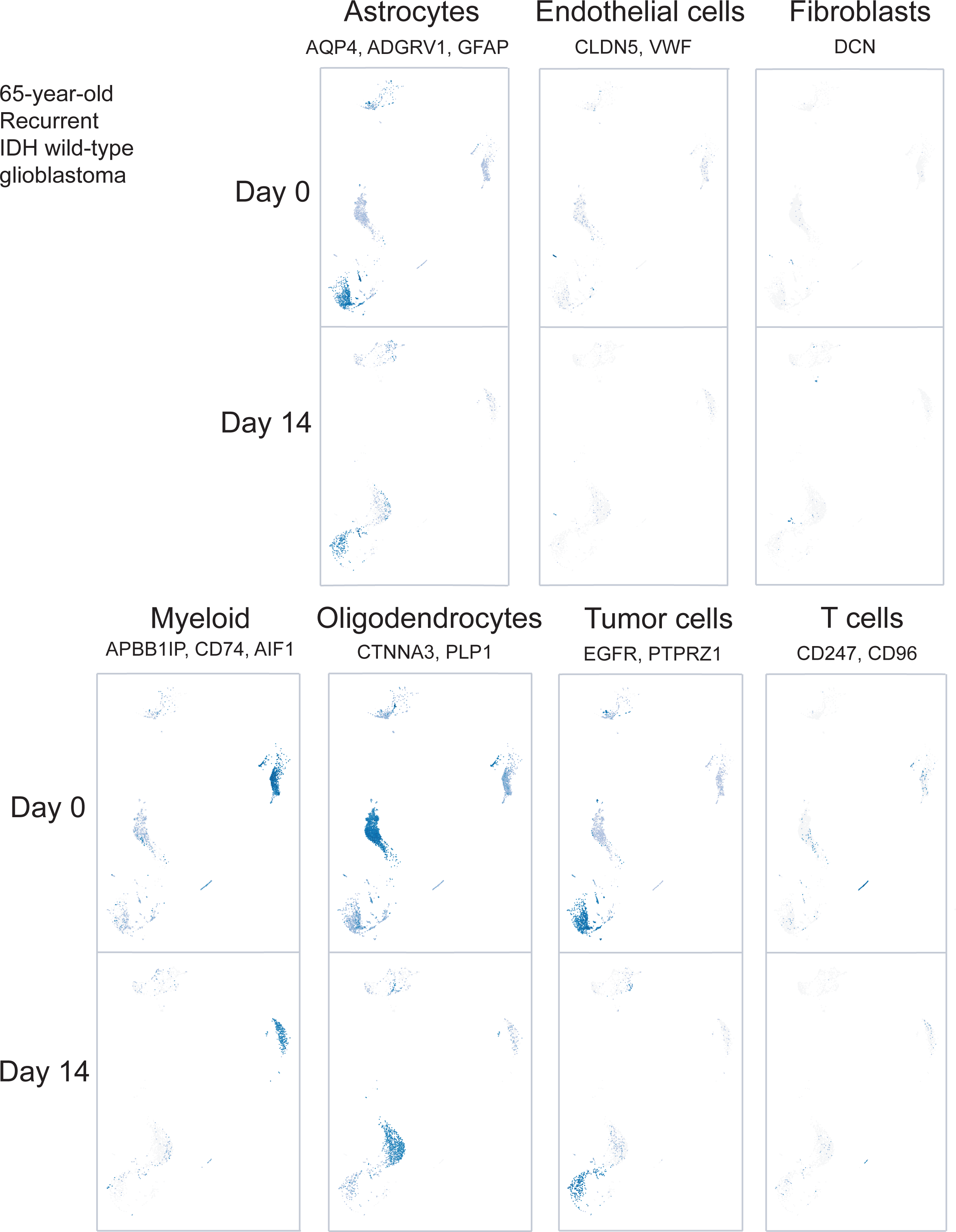
Marker genes used to assign cell types for tissue E. In cases where multiple marker genes were used, feature sum was used across all the genes listed.

**Supplemental Figure 6.**
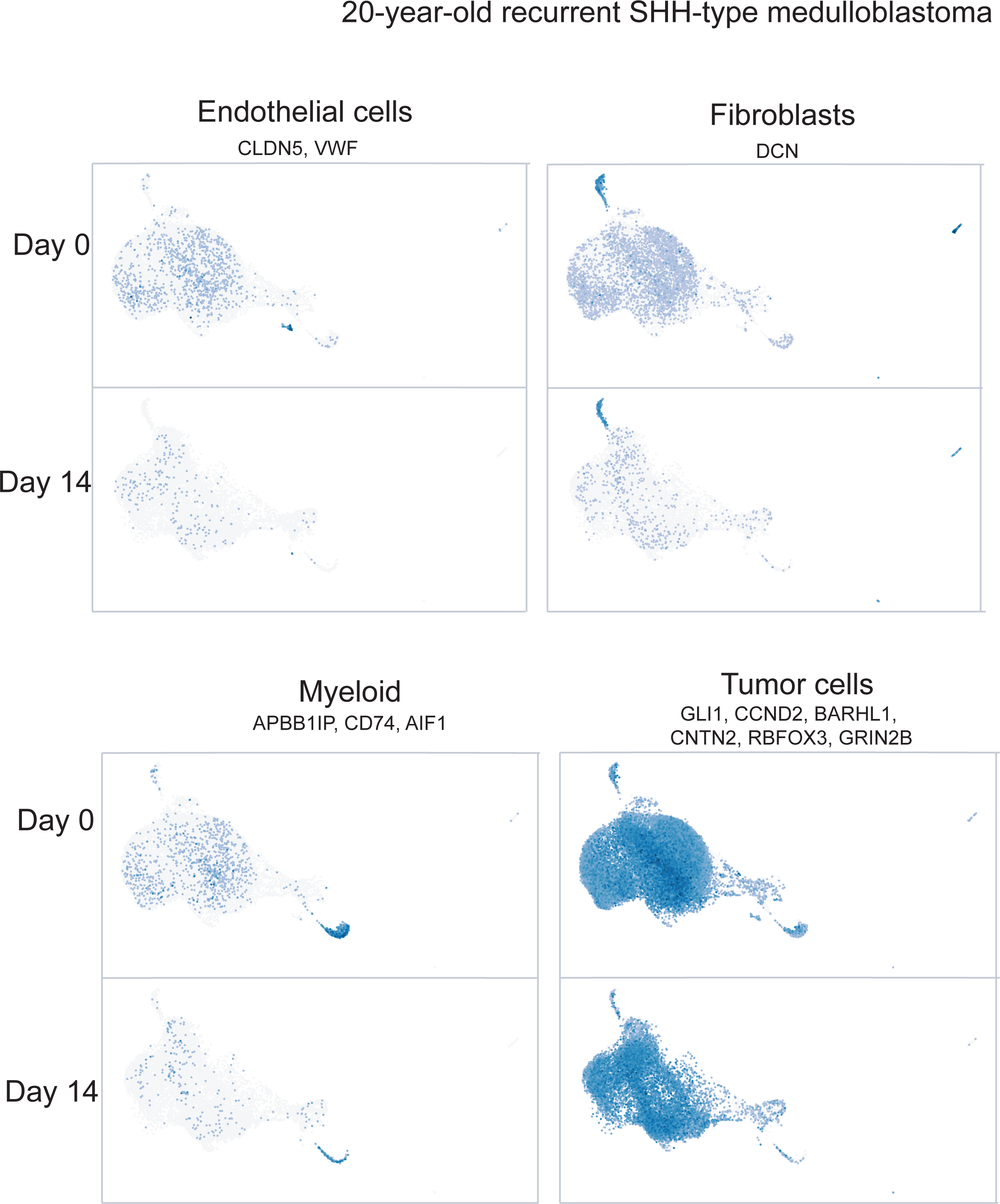
Marker genes used to assign cell types for tissue F. In cases where multiple marker genes were used, feature sum was used across all the genes listed.

**Supplemental Figure 7.**
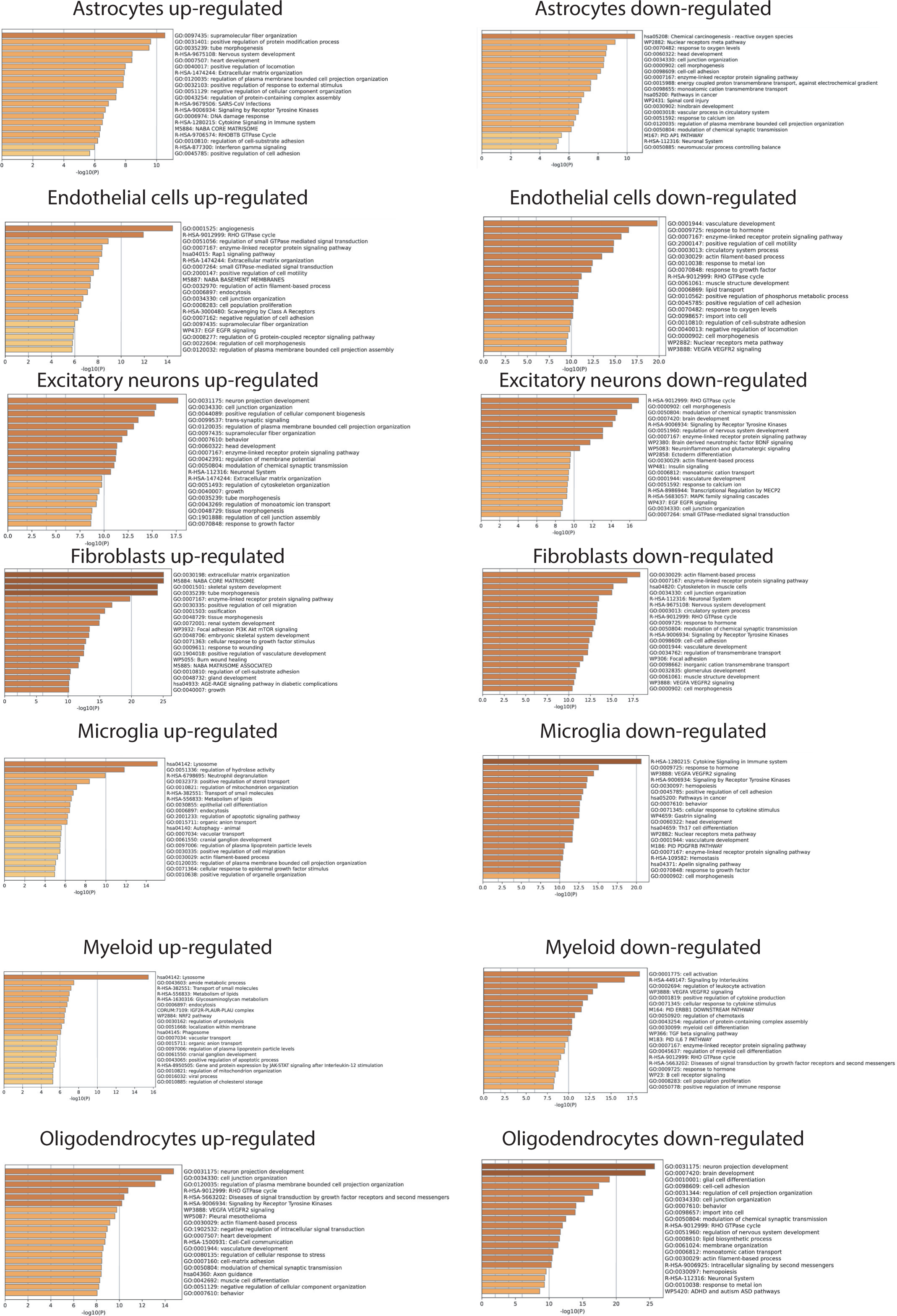
Gene ontology analysis for all non-neoplastic cell types. The genes showing significant up- or down-regulation in at least two patient samples were compiled and entered into Metascape, a web browser tool that generates gene ontology, cell pathway, and disease features. In none of the cells was a coherent transcriptional shift seen.

**Supplemental Figure 8.**
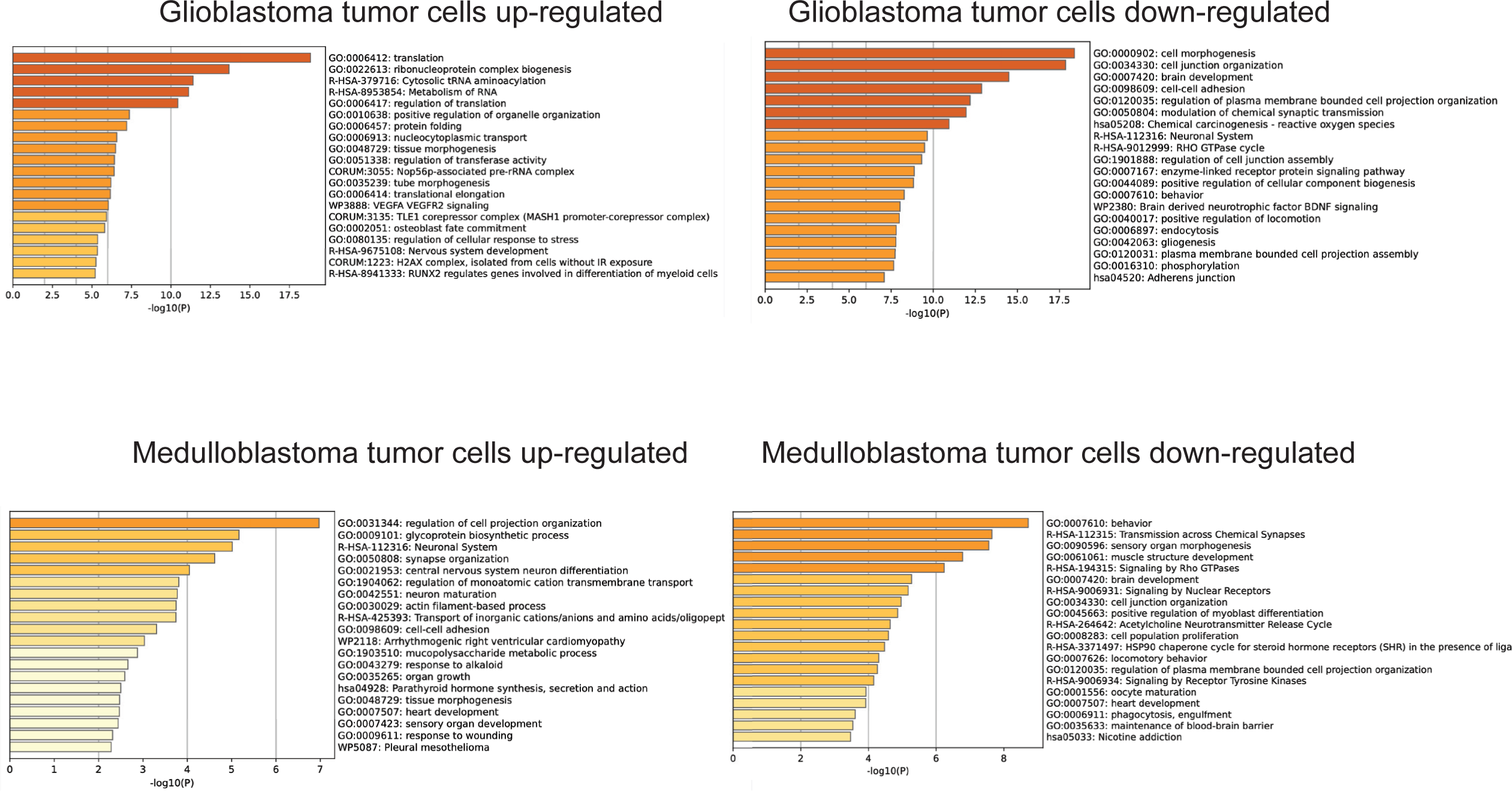
Gene ontology analysis for all neoplastic cell types. The genes showing significant up- or down-regulation in at least two patient samples were compiled and entered into Metascape, a web browser tool that generates gene ontology, cell pathway, and disease features.

**Supplemental Table 1. Up- and down-regulated genes over fourteen days in culture by cell type and tissue in which the genes were identified.**

## References

1. Loewa A, Feng JJ, Hedtrich S. Human disease models in drug development. Nat Rev Bioeng. 2023 May 11:1–15.

2. Scannell JW, Bosley J. When Quality Beats Quantity: Decision Theory, Drug Discovery, and the Reproducibility Crisis. PLoS One. 2016 Feb 10;11(2):e0147215.

3. He Z, Dony L, Fleck JS, Szałata A, Li KX, Slišković I, Lin HC, Santel M, Atamian A, Quadrato G, Sun J, Pașca SP; Human Cell Atlas Organoid Biological Network; Camp JG, Theis FJ, Treutlein B. An integrated transcriptomic cell atlas of human neural organoids. Nature. 2024 Nov;635(8039):690–698. doi: 10.1038/s41586-024-08172-8. Epub 2024 Nov 20. Erratum in: Nature. 2024 Dec 11.

4. Chaichana KL, Capilla-Gonzalez V, Gonzalez-Perez O, Pradilla G, Han J, Olivi A, Brem H, Garcia-Verdugo JM, Quiñones-Hinojosa A. Preservation of glial cytoarchitecture from ex vivo human tumor and non-tumor cerebral cortical explants: A human model to study neurological diseases. J Neurosci Methods. 2007 Aug 30;164(2):261–70.

5. Schwarz N, Hedrich UBS, Schwarz H, P A H, Dammeier N, Auffenberg E, Bedogni F, Honegger JB, Lerche H, Wuttke TV, Koch H. Human Cerebrospinal fluid promotes long-term neuronal viability and network function in human neocortical organotypic brain slice cultures. Sci Rep. 2017 Sep 25;7(1):12249.

6. Ting JT, Lee BR, Chong P, Soler-Llavina G, Cobbs C, Koch C, Zeng H, Lein E. Preparation of Acute Brain Slices Using an Optimized N-Methyl-D-glucamine Protective Recovery Method. J Vis Exp. 2018 Feb 26;(132):53825.

7. Ting JT, Kalmbach B, Chong P, de Frates R, Keene CD, Gwinn RP, Cobbs C, Ko AL, Ojemann JG, Ellenbogen RG, Koch C, Lein E. A robust ex vivo experimental platform for molecular-genetic dissection of adult human neocortical cell types and circuits. Sci Rep. 2018 May 30;8(1):8407.

8. Park TI, Smyth LCD, Aalderink M, Woolf ZR, Rustenhoven J, Lee K, Jansson D, Smith A, Feng S, Correia J, Heppner P, Schweder P, Mee E, Dragunow M. Routine culture and study of adult human brain cells from neurosurgical specimens. Nat Protoc. 2022 Feb;17(2):190–221.

9. Bak A, Koch H, van Loo KMJ, Schmied K, Gittel B, Weber Y, Ort J, Schwarz N, Tauber SC, Wuttke TV, Delev D. Human organotypic brain slice cultures: a detailed and improved protocol for preparation and long-term maintenance. J Neurosci Methods. 2024 Apr;404:110055.

10. Andrews JP, Geng J, Voitiuk K, Elliott MAT, Shin D, Robbins A, Spaeth A, Wang A, Li L, Solis D, Keefe MG, Sevetson JL, Rivera de Jesús JA, Donohue KC, Larson HH, Ehrlich D, Auguste KI, Salama S, Sohal V, Sharf T, Haussler D, Cadwell CR, Schaffer DV, Chang EF, Teodorescu M, Nowakowski TJ. Multimodal evaluation of network activity and optogenetic interventions in human hippocampal slices. Nat Neurosci. 2024 Dec;27(12):2487–2499.

11. Allen Institute for Brain Science. Cell types database: RNA-seq data from human M1 10x. Available at: https://celltypes.brain-map.org/rnaseq/human_m1_10x?selectedVisualization=Heatmap&colorByFeature=Cell+Type&colorByFeatureValue=GAD1. Accessed repeatedly Jan 2024 – Nov 2024.

12. Winkler EA, Kim CN, Ross JM, Garcia JH, Gil E, Oh I, Chen LQ, Wu D, Catapano JS, Raygor K, Narsinh K, Kim H, Weinsheimer S, Cooke DL, Walcott BP, Lawton MT, Gupta N, Zlokovic BV, Chang EF, Abla AA, Lim DA, Nowakowski TJ. A single-cell atlas of the normal and malformed human brain vasculature. Science. 2022 Mar 4;375(6584):eabi7377.

13. Abdelfattah N, Kumar P, Wang C, Leu JS, Flynn WF, Gao R, Baskin DS, Pichumani K, Ijare OB, Wood SL, Powell SZ, Haviland DL, Parker Kerrigan BC, Lang FF, Prabhu SS, Huntoon KM, Jiang W, Kim BYS, George J, Yun K. Single-cell analysis of human glioma and immune cells identifies S100A4 as an immunotherapy target. Nat Commun. 2022 Feb 9;13(1):767.

14. Wang X, Sun Q, Wang W, Liu B, Gu Y, Chen L. Decoding key cell sub-populations and molecular alterations in glioblastoma at recurrence by single-cell analysis. Acta Neuropathol Commun. 2023 Jul 31;11(1):125.

15. Ocasio JK, Babcock B, Malawsky D, Weir SJ, Loo L, Simon JM, Zylka MJ, Hwang D, Dismuke T, Sokolsky M, Rosen EP, Vibhakar R, Zhang J, Saulnier O, Vladoiu M, El-Hamamy I, Stein LD, Taylor MD, Smith KS, Northcott PA, Colaneri A, Wilhelmsen K, Gershon TR. scRNA-seq in medulloblastoma shows cellular heterogeneity and lineage expansion support resistance to SHH inhibitor therapy. Nat Commun. 2019 Dec 20;10(1):5829. Erratum in: Nat Commun. 2022 May 26;13(1):3048.

16. Zhou Y, Zhou B, Pache L, Chang M, Khodabakhshi AH, Tanaseichuk O, Benner C, Chanda SK. Metascape provides a biologist-oriented resource for the analysis of systems-level datasets. Nat Commun. 2019 Apr 3;10(1):1523.

17. Iwasaki Y, Bernou C, Gorda B, Colomb S, Ganesh G, Gaudin R. Organotypic culture of post-mortem adult human brain explants exhibits synaptic plasticity. Brain Stimul. 2024 Sep-Oct;17(5):1018–1023.

18. Schwanhäusser B, Busse D, Li N, Dittmar G, Schuchhardt J, Wolf J, Chen W, Selbach M. Global quantification of mammalian gene expression control. Nature. 2011 May 19;473(7347):337–42. Erratum in: Nature. 2013 Mar 07;495(7439):126-7.

